# Electrostatic interactions control the adsorption of extracellular vesicles onto supported lipid bilayers

**DOI:** 10.1101/2023.04.14.536633

**Authors:** Andrea Ridolfi, Jacopo Cardellini, Fatlinda Gashi, Martijn J.C. van Herwijnen, Martin Trulsson, José Campos-Terán, Marca H. M. Wauben, Debora Berti, Tommy Nylander, Joakim Stenhammar

## Abstract

Communication between cells located in different parts of an organism is often mediated by membrane-enveloped nanoparticles, such as extracellular vesicles (EVs). EV binding and cell uptake mechanisms depend on the heterogeneous composition of the EV membrane. From a colloidal perspective, the EV membrane interacts with other biological interfaces via both specific and non-specific interactions, where the latter include long-ranged electrostatic and van der Waals forces, and short-ranged repulsive “steric-hydration” forces. While electrostatic forces are generally exploited in most EV immobilization protocols, the roles played by various colloidal forces in controlling EV adsorption on surfaces have not yet been thoroughly addressed. In the present work, we study the interaction and adsorption of EVs with supported lipid bilayers (SLBs) carrying different surface charge densities. By probing the EV-SLB interaction using quartz crystal microbalance with dissipation monitoring (QCM-D) and confocal laser scanning microscopy (CLSM), we demonstrate that EV adsorption onto lipid membranes can be controlled by varying the strength of electrostatic forces. We then model the observed phenomena within the framework of nonlinear Poisson-Boltzmann theory. Modelling results confirm the experimental observations and highlight the crucial role played by attractive electrostatics in EV adsorption onto lipid membranes. Our results provide new fundamental insights into EV-membrane interactions and could be useful for developing novel EV separation and immobilization strategies.

## Introduction

Cell-to-cell communication is involved in many biological processes and in the onset and spread of multiple pathological conditions in multicellular organisms. Depending on the location, relative distance, and type of message to be delivered, cells can use several different communication pathways^1,2^. One of the most powerful strategies for long-distance communication between cells located in different parts of an organism is the production and secretion of membrane-enveloped nanoparticles^3,4^, where one of the most intriguing classes is extracellular vesicles (EVs)^5–7^. EVs are unilamellar vesicles, typically in the nanometric range, used by cells as carriers for biological material such as proteins, carbohydrates and nucleic acids^8,9^. EV binding and fusion with, or uptake by, target cells occur via multiple different mechanisms. Many of them, such as vesicle fusion and raft- and receptor-mediated endocytosis, strongly depend on the heterogeneous composition of the EV membrane^10^, which consists of a multicomponent lipid bilayer containing a wide range of membrane proteins and biomolecules^8^.

From a colloidal perspective, the EV membrane can interact with other biological interfaces *via* both specific and non-specific interactions^3,11,12^. The latter category is typically dominated by electric double layer and van der Waals forces, as outlined in the DLVO theory of colloidal stability, and short-ranged repulsive “steric-hydration” forces of unclear molecular origin that have been measured between hydrated lipid bilayers^13,14^. Several studies have demonstrated that phenomena such as adsorption, deformation, and rupture of synthetic lipid vesicles in contact with charged substrates are strongly dependent on the magnitude of electrostatic interactions^15–17^. Moreover, electrostatic interactions are commonly exploited to adsorb and immobilize EVs and synthetic lipid vesicles on solid substrates to study their properties^18–21^., The most common setup to study the interaction of lipid vesicles with lipid membranes under controlled and simplified conditions consists of a supported lipid bilayer (SLB), *i*.*e*., a planar lipid bilayer deposited onto a rigid support^22–25^. Recent investigations have specifically probed the interaction of biologically derived EVs with SLBs^26,27^, however without addressing the roles of generic colloidal forces in controlling EV adsorption.

In the present work, we employ weakly negatively charged EVs derived from bovine milk as a model system to study EV adsorption on synthetic SLBs with varying positive surface charge densities. This setup enables us to study the specific roles played by electrostatics in the interactions between EVs and other lipid interfaces, thus avoiding the compositional and topological complexity associated with natural membranes. To generate charge-controlled SLBs, we use mixed liposomes composed of zwitterionic 1,2-dioleoyl-sn-glycero-3-phosphocholine (DOPC) and cationic 1,2-dioleoyl-3-trimethylammonium-propane (DOTAP). Tuning the ratio between the two amphiphilic molecules allows us to accurately control the surface charge density of the SLBs. We quantified the adsorption of EVs on the SLBs using two independent experimental techniques: quartz crystal microbalance with dissipation monitoring (QCM-D) and confocal laser scanning microscopy (CLSM). The two characterization techniques independently demonstrate that EV adsorption onto lipid membranes can be readily controlled by varying the magnitude of the positive surface charge density of the SLB. For SLBs composed of less than 40% w/w of DOTAP, corresponding to low surface charge densities, EV adsorption is negligible. In contrast, for higher contents of cationic DOTAP molecules EV adsorption increases monotonically with the DOTAP/DOPC ratio. We next rationalized the EV-SLB interactions within the framework of nonlinear Poisson-Boltzmann (PB) theory, which accurately describes the electrostatic interactions between unequally charged surfaces. When the resulting interaction free energy curves were combined with empirically measured short-range forces known to act between lipid bilayers, they confirmed that attractive electrostatic interactions are indeed expected to be the dominant driver of EV adsorption for the experimentally probed DOTAP/DOPC ratios.

## Materials and Methods

### Liposome preparation for QCM experiments

1,2-dioleoyl-sn-glycero-3-phosphocholine (DOPC) and 1,2-dioleoyl-3-trimethylammonium-propane (DOTAP) dry powders were purchased from Sigma Aldrich (St. Louis, MO, USA). Different amounts of DOPC and DOTAP were used to tune the surface charge of the employed liposomes; Table S1 reports the DOTAP/DOPC w/w percentages employed in the study, together with the nomenclature used in the next sections to describe them. In general, to prepare liposomes, the amount of DOPC and DOTAP powders corresponding to the desired final composition were weighed in to yield a total mass of 40 mg, before being dissolved in chloroform. Chloroform was then gently evaporated using a stream of nitrogen, leaving a thin lipid film at the bottom of the vials. The vials were covered with aluminum foil and put into a vacuum chamber overnight to allow the lipid films to dry completely. The following day, the lipid films were hydrated using 4 ml of PBS and the dispersions were successively extruded using 200 nm NanoSizer MINI Liposome Extruder (T&T Scientific). The extruded liposome dispersions were further diluted with PBS to a final concentration of 0.25 mg/ml for the following experiments.

### EV isolation and purification

Raw bovine milk (100 ml) was collected from a cooled tank in a local dairy farm (Tolakker, Utrecht, The Netherlands), transferred to 50 ml polypropylene tubes and centrifuged for 10 minutes at 22°C at 3000 xg (Beckman Coulter Allegra X-12R, Fullerton, CA, USA). After removal of the cream layer, the milk supernatant was harvested without disturbing the pellet and transferred to new tubes. A second centrifugation step at 3000 xg followed, after which the milk supernatant was collected and stored at −80°C until further processing. To isolate milk EVs, thawed milk supernatant was transferred to polyallomer SW40 tubes (Beckman Coulter) and centrifuged at 5000 xg for 30 minutes at 4°C and subsequently at 10000 xg (Beckman Coulter Optima L-90K with a SW40Ti rotor). For the precipitation of caseins, the milk supernatant was acidified to pH 4.6 by adding hydrochloric acid (HCl, 1M) while stirring. Caseins were pelleted by centrifugation at 360 xg (Beckman Coulter Allegra X-12R) for 10 minutes at 4°C, after which casein-free milk supernatant was collected and neutralized to pH 7.0 with sodium hydroxide (NaOH 1M). Next, 6.5 ml of the milk 10000 xg supernatant was loaded on top of a 60% – 10% Optiprep gradient (OptiprepTM, Progen Biotechnik GmbH, Heidelberg, Germany) made in a SW40 tube. Gradients were ultracentrifuged at 192,000 xg (Beckman Coulter Optima L-90K with a SW40Ti rotor) for 15 − 18 h. After centrifugation, fractions of 500 μl were harvested and densities were measured to identify the EV-containing fractions (Density 1.06 − 1.19 g/ml), which were pooled. Optiprep was removed during size exclusion chromatography of the pooled EV-containing fractions using a 20 ml column (Bio-Rad Laboratories, Hercules, CA, USA) packed with 15 ml Sephadex g100 (Sigma-Aldrich, St. Louis, MO, USA). Fractions of 1 ml were eluted from the column with PBS (GibcoTM, Invitrogen, Carlsbad, CA, USA). The EV-containing fractions 3 to 9 were pooled and stored at −80°C until use. For all the adsorption experiments (QCM-D and CLSM), milk EV preparations were diluted 1:200. The procedure of bovine milk EV isolation, as well as the analysis of milk EV preparations by Colorimetric Nanoplasmonic Assay (CONAN), Atomic Force Microscopy (AFM) and by Western-blotting has been previously described^21^ and reported in the EV-TRACK knowledgebase (EV190077).

### DLS and ζ potential measurements

Dynamic light scattering (DLS) measurements were performed using a Zetasizer Nano ZS (Malvern Pananalytical Ltd., Malvern, UK), yielding information about the size distribution of both liposomes and EVs. Measurements were performed in triplicate, each one comprising 7 runs of 30 seconds each. The same instrument was then used for measuring the *ζ* potential of the lipid vesicles; measurements were performed in triplicate and comprised 10 runs with automatic attenuation and optical settings for all samples. For both DLS and *ζ* potential, the measurements were performed in PBS at a constant temperature of 25°C. The hydrodynamic radii of the vesicles are reported in Table S2, together with the respective autocorrelation functions and fits obtained from DLS analysis (Figure S1). The obtained results and error values are expressed as the average and standard deviation of the respective triplicate for each sample.

### Nanoparticle tracking analysis

Purified bovine milk EV preparations were quantified by nanoparticle tracking analysis (NTA) according to the procedure described previously^28^, employing a NanoSight NS300 instrument (Malvern Instruments Ltd, UK) equipped with a 405 nm laser and a high sensitivity sCMOS camera. The NanoSight NTA software version 3.2.16 was used for data acquisition and processing. The EV sample was diluted in endotoxin-free Dulbecco’s 1X PBS without Ca^2+^ and Mg^2+^ (TMS -012-A Millipore). Three technical replicates were measured in various dilutions ranging from 1:500 to 1:1000 in a 60-second video under controlled temperature set at 20*°*C.

### QCM-D experiments

Quartz crystal microbalance with dissipation monitoring (QCM-D) experiments were performed using a Q-Sense E4 instrument (Biolin Scientific, Göteborg, Sweden) using silicon oxide coated 5 MHz AT-cut quartz crystal sensor chips (Biolin Scientific) as substrate for the vesicle deposition. Prior to each experiment, the sensor chips were sequentially immersed in Hellmanex 2%, Ethanol and MilliQ water under sonication in an ultrasonic bath for 7 minutes for each solvent. After being dried in a stream of nitrogen, the sensor chips were then further cleaned in an air plasma for 1 min at 0.02 mbar using a plasma cleaner (Harrick Scientific Corp, model PDC-3XG, New York, USA). The sensor chips were then immediately placed in the sealed QCM-D measurement chambers and buffer was flowed through the chamber. The temperature was fixed at 25°C and kept constant throughout all the experiments. During the QCM-D measurements, variations in both the resonance frequency and energy dissipation of the sensor chips were recorded as a function of time. To form the SLBs, liposomes were flushed within the QCM-D measurement chamber, where they adsorbed on the sensor surface. Then, MilliQ water was flushed to create an osmotic imbalance and promote the rupture of still intact liposomes. Once the whole sensor surface was covered with SLBs, indicated by a frequency drop of ∼20 − 30 Hz and negligible change in dissipation, the buffer was changed to PBS and the systems were left to equilibrate for approximately 1 hour. After that, EVs were injected at a flow rate of 0.8 ml/min. The outlet solution was collected and re-injected within the QCM-D chamber three times, to maximize the number of EVs interacting with the SLBs. The flow was then stopped and the systems were left to equilibrate for approximately 1 hour before recording the frequency shift values. The results reported in the manuscript were obtained by taking the average of the values recorded from the 5^th^, 7^th^, 9^th^ and 11^th^ harmonics at specific times (see Figure 1), while the error bars refer to the standard deviation of the four harmonics.

**Figure 1:**
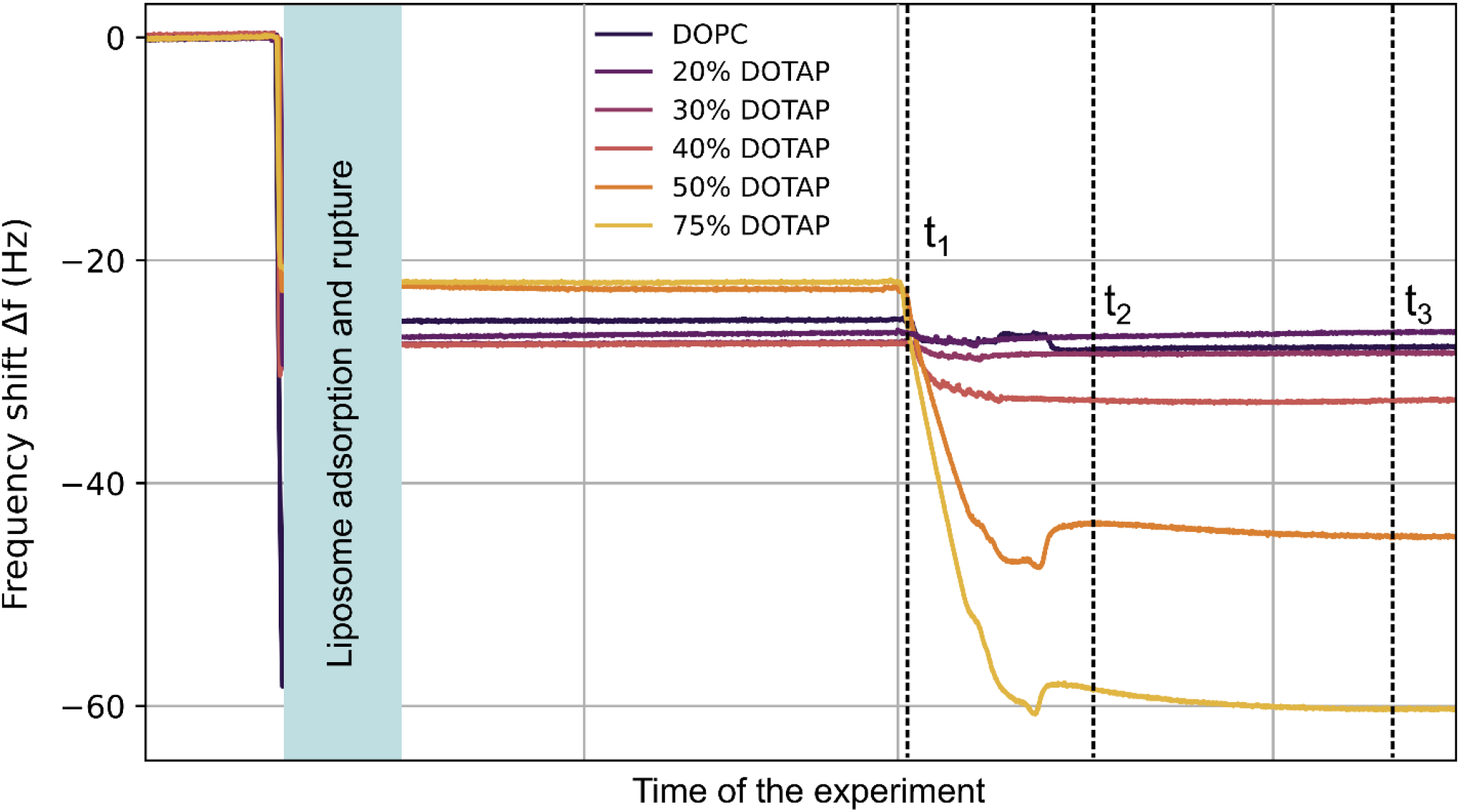
Representative frequency shifts Δ*f* corresponding to the 5^th^ harmonic for the SLB-EV systems with various SLB compositions, as indicated. The different regimes correspond to the starting baseline (0 Hz), the SLB formation (first plateau), the EV injection (*t*_1_), adsorption and equilibration of EVs on the SLB (*t*_2_ and *t*_3_, respectively).

### CLSM experiments

For the Confocal Laser Scanning Microscopy (CLSM) experiments, liposomes and EVs were respectively labelled with two different fluorescent probes (0.1 mol%), characterized by well-separated emission spectra. Fluorescent liposomes and EVs were obtained by incubating the vesicle dispersions overnight with β-Bodipy TM FL C12-HPLC (emission wavelength 512 nm, Thermofisher) and 18:1 Cyanine-5-Phosphatidylethanolamine (Cy5, emission wavelength 663 nm, Avanti Polar Lipids) dry films, respectively. The SLBs were obtained by depositing 100 μL of a 0.1 M NaCl dispersion containing the respective liposomes (at a lipid concentration of 0.5 mg/ml) in three different Borosilicate CLSM wells. Vesicle rupture and SLB formation were then achieved via osmotic shock by gently rinsing the sample with Milli-Q water. To avoid the presence of intact vesicles in the wells, each sample was further rinsed 15 times with MilliQ water. Finally, EVs were injected into the wells from the top using a micropipette. A Leica CLSM TCS SP8 confocal microscope was used to probe the EV-SLB interaction. The fluorescent β-Bodipy probe was excited at 488 nm and its emission was collected in the range 498 − 530 nm with a photomultiplier tube (PMT), while Cy5 was excited at a wavelength of 633 nm and collected in the 650 − 700 nm range. Images were taken at a resolution of 512 × 512 pixels, after leaving the systems (SLBs or SLBs with EVs) equilibrating for approximately 1 hour. Leica Application Suite X (LAS X) software was used to create three-dimensional reconstructions of the *z*-stacks.

### Theoretical modelling

As is further discussed in the Results and Discussion section, we numerically solved the PB equation (4) for two planar surfaces separated by a distance *D* under the assumption of constant surface charge density, which is typically an accurate choice for colloidal systems that do not possess a strong propensity to charge regulate through titration of surface groups or adsorption of charged species. This leads to the boundary conditions 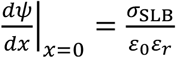 and 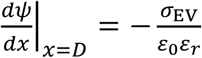, where *ψ*(*x*) is the electrostatic potential, *σ*_SLB_ and *σ*_EV_ are the SLB and EV surface charge densities, and *ε*_0_*ε*_*r*_ is the solvent permittivity. For the SLB surface charge density, we used the estimated values of *σ*_SLB_ shown in Table 1, calculated using Eq. (3). For the EVs, the surface charge density was adjusted to fit measured adsorption data from CLSM, as described in Results and Discussion. We numerically integrated the PB equation to give the potential profile *ψ*(*x*), which can be directly translated into the osmotic pressure, Π, between the two planar surfaces according to^29^

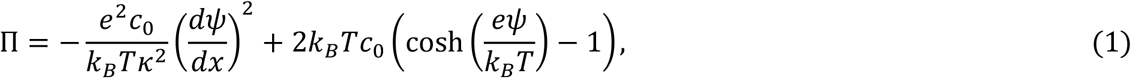

where *e* is the elementary charge, κ the inverse Debye screening length, *c*_0_ = 150 mM the bulk salt concentration, and *k*_*B*_*T* the thermal energy, where we used *T* = 298 K. It should be noted that the right-hand-side is independent of *x*, since the osmotic pressure is uniform across the gap. Integrating the osmotic pressure as a function of the gap distance *D* then yields an expression of the interaction free energy per unit area *A*_PP_(*D*) between the two planes. We then used the Derjaguin approximation^13^ to account for the spherical shape of the EV, yielding the sphere-plane force

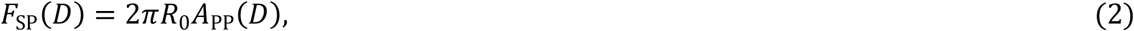

Where *R*_0_ is the EV radius, which we set equal to the average hydrodynamic radius of 94.5 nm, measured for the EV sample from the DLS experiments. Finally, numerically integrating the force *F*_SP_ with respect to *D* once more yields the electrostatic interaction free energy *A*_EI_(*D*) between the planar SLB and spherical EV.

**Table 1:** Lipid compositions, measured *ζ* potential and expected SLB surface charge densities calculated from the lipid compositions of the studied model liposomes.

| Liposome composition | Measured $\zeta$ potential (mV) | Calculated surface charge density $\sigma$ (C/m<sup>2</sup>) |
| --- | --- | --- |
| Pure DOPC | $-0.9 \pm 0.3$ | 0.00 |
| DOTAP 20% | $18 \pm 1$ | 0.051 |
| DOTAP 30% | $24 \pm 2$ | 0.077 |
| DOTAP 40% | $27 \pm 1$ | 0.103 |
| DOTAP 50% | $30 \pm 1$ | 0.130 |
| DOTAP 75% | $32.1 \pm 0.9$ | 0.197 |
| EVs | $-7.7 \pm 0.9$ | -0.0094* |

\* $\sigma_{\text{EV}}$ was adjusted to match the measured EV adsorption as described in Results and Discussion.

## Results and Discussion

### Characterization of EV and SLB electrostatic properties

Due to their composition, EVs generally possess a weakly negative *ζ* potential, indicating a net negative surface charge density of their lipid membrane^30^. In our milk EV sample, the measured *ζ* potential was −77 ± 0.8 mV. DLS measurements showed a relatively monodisperse size distribution, with an average vesicle hydrodynamic radius of *R*_0_ = 95 ± 1 nm and a polydispersity index (PDI) of 0.218 ± 0.029. The *ζ* potential and size measurements were performed in PBS buffer, corresponding to an inverse Debye screening length of κ = 1.26 · 10^9^ m^-1^. Nanoparticle Tracking Analysis (NTA) was used to measure the vesicle size and concentration, yielding an average vesicle radius of 76 ± 3 nm and an EV concentration of 1.84 × 10^11^ particles per ml (Figure S2). The discrepancy in the EV radii measured from NTA and DLS is because the latter technique is biased towards larger particles, since they scatter light more intensely. For further experimental details, we refer to Materials and Methods and Supplementary Information (SI).

The SLBs were prepared with different DOPC/DOTAP ratios to reveal how electrostatics influences vesicle-membrane interactions. Since DOTAP is positively charged, we obtained synthetic membranes with a controllable surface charge density. Table 1 reports all the liposome compositions used to produce the studied SLBs. As expected, the magnitude of the liposome *ζ* potential increases with the concentration of DOTAP. The theoretical SLB surface charge density *σ*_SLB_, also presented in Table 1, was then calculated from the lipid composition and the approximate headgroup areas of the two lipid species, assuming that each DOTAP molecule carries a single elementary charge independently of the membrane composition. The latter is a reasonable assumption given that the charge on DOTAP is not titratable; therefore, we do not expect the SLBs to possess a strong ability to regulate charge in response to changes in its electrostatic environment, *i*.*e*., the lipid composition or the ionic strength. The resulting expression is given by

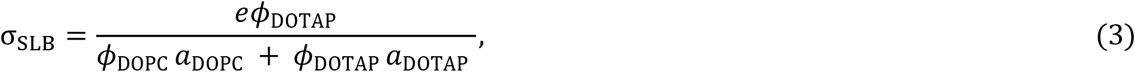

where *ϕ*_DOPC_ and *ϕ*_DOTAP_ are the percentage fractions of the respective lipid constituents, *e* is the electron charge, and *a*_DOPC_ and *a*_DOTAP_ the headgroup areas of the lipid molecules, taken to be 63.3 Å^2^ and 60.4 Å^2^, respectively^31^. Since DOPC and DOTAP are miscible and form homogeneous bilayers for most of the probed concentrations^32^, we treat the formed SLBs as homogeneously charged surfaces with uniform surface charge density *σ*_SLB_. The size range and polydispersity of the liposomes, as measured by Dynamic Light Scattering (DLS) are presented in SI and were furthermore essentially found to be independent of the lipid composition.

### Measurement of EV adsorption through QCM-D

We first used QCM-D to study the adsorption of EVs onto SLBs of varying surface charge density, as reported in Table 1. To form the SLBs, liposomes with different DOPC/DOTAP ratios were injected into the QCM-D flow chamber. Due to mutual interaction and crowding on the sensor surface, the liposomes eventually ruptured and fused, forming uniform SLBs. To promote the rupture of still intact vesicles, milliQ water was flushed through the cell in a second step to create the osmotic imbalance needed to destabilize and rupture the remaining vesicles. The successful formation of continuous SLBs uniformly covering the entire sensor surface is confirmed by the frequency shift (Δ*f*) values displayed in Figure 1, ranging from ∼20 to ∼30 Hz^33^, together with the negligible shifts in the dissipation signals (Δ*D*) relative to the clean crystal (SI)^34,35^. Since the differently charged liposomes adsorbed and formed SLBs according to different mechanisms (*e*.*g*., adsorption to the substrate, vesicle rupture and fusion) and time frames, the portions of the plots describing these phenomena have been omitted from Fig. 1 in order to better compare the differences in the interactions between EV and the various SLBs; the full QCM-D plots including the adsorption and rupture phases are reported in Fig. S3. The small differences between the Δ*f* values at the first plateau, corresponding to the point when the equilibrated SLBs covered the entire sensor surface, can be attributed to both their differing compositions and the variability between different experiments.

After SLB equilibration, EVs were injected into the flow chamber at a flow rate of 0.8 ml/min, indicated by the point *t*_1_ in Fig. 1. The sudden drop in Δ*f* indicates a significant interaction of the EVs with the SLB surface. The extent of the Δ*f* drop strongly depends on the surface charge density of the SLBs: while the pure DOPC SLB, formed from liposomes that are characterized by a weakly negative *ζ* potential, displays a negligible frequency drop, the highly positively charged 75% DOTAP SLB features the highest Δ*f*, indicating significant EV adsorption. After the injection phase, at the point marked by *t*_2_ in Fig. 1, the flow was stopped and the EVs were left to equilibrate with the SLBs for one hour until a new stable plateau in Δ*f* was reached (point *t*_3_ in Fig. 1). The Δ*f* values recorded for the EV adsorption at steady state, at point *t*_3_, were then used for evaluating the EV adsorption at equilibrium. The bar plot in Fig. 2 reports the difference in the Δ*f* and Δ*D* values between *t*_3_ (after equilibration) and *t*_1_ (before EV injection) for the different SLBs, which indicates of the amount of EVs adsorbed onto the respective SLBs. The results confirm that EV adsorption strongly depends on the SLB surface charge density: for SLBs with DOTAP fractions lower than 40%, EV adsorption is negligible with no significant difference compared to the pure DOPC bilayer. EV adsorption starts to occur for SLBs with a DOTAP percentage of 40% and then increases sharply with the concentration of the cationic phospholipid. After EV adsorption, the dissipation signals increase proportionally to the DOTAP concentration, eventually reaching Δ*D* values in the 9 − 11.5 ppm range (Figs. 2 and S3); such large dissipation shifts suggest the formation of a layer of intact EVs oscillating out of phase with respect to the underlying SLB. This interpretation is further strengthened by the dependency of both Δ*f* and Δ*D* on the DOTAP concentration and by the spreading of the different harmonics (Fig. S3). If instead EVs had ruptured and merged with the SLB, the frequency and dissipation shifts should be similar for all membrane compositions where adsorption occurs. More specifically, Δ*f* would plateau at a constant value and Δ*D* would be close to zero, with only small differences between the different harmonics. Even though recent works^26,27^ have demonstrated EV-SLB fusion after adsorption, these have relied on more specific interactions, respectively the adsorption to the boundaries between liquid-ordered and liquid-disordered phases^26^ and antibody-mediated attractive interactions^27^. In contrast, we expect our system to lack any strong attractive interactions apart from generic electrostatic and van der Waals interactions, which have previously been shown insufficient to induce membrane fusion^36^.

**Figure 2:**
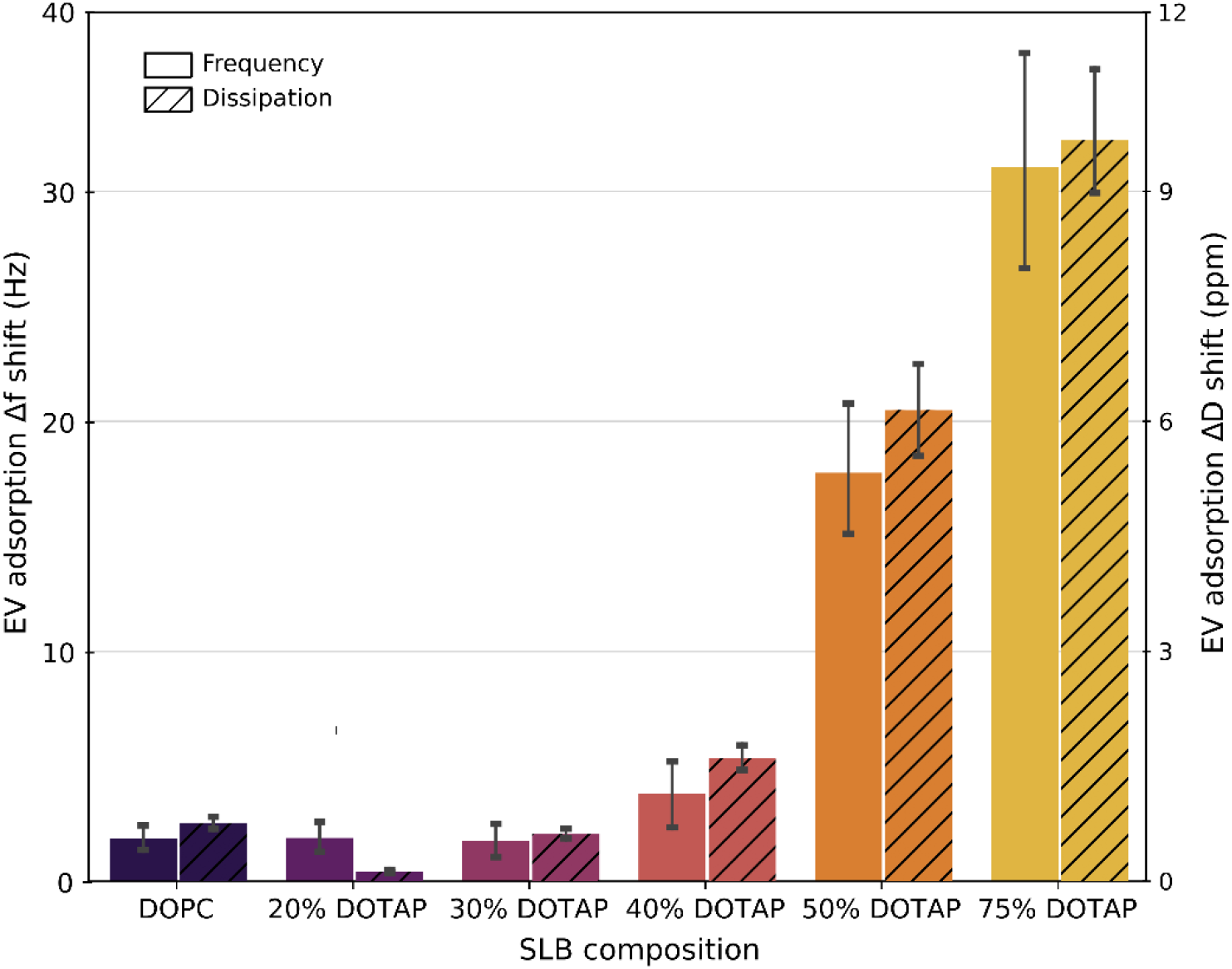
Average QCM-D frequency and dissipation shifts (Δ*f* and Δ*D*, respectively) due to EV adsorption on the various SLBs, as indicated. The shifts are measured as the difference between the Δ*f* or Δ*D* values recorded at times *t*_*3*_ and *t*_*1*_ illustrated in Fig. 1. The plotted values represent an average over the 5^th^, 7^th^, 9^th^, and 11^th^ harmonics, and error bars indicate the standard deviation of the four values.

Extending this qualitative interpretation of QCM-D results to properly account for the formation of multilayered viscoelastic systems, such as vesicles adsorbed on top of soft SLBs, is however highly complex. Specifically, translating the obtained Δ*f* values into more physical parameters such as adsorbed mass is not straightforward. For example, the results are influenced by dissipative phenomena related to the water trapped within the vesicles and between the multiple layers^37,38^. While the Voigt model^37^ could in principle be used for modeling the adsorption of such viscoelastic soft layers, this model requires *a priori* knowledge of properties such as the density and viscosity of the adsorbed layers, which in this case are both unknown and highly dependent on the morphology of the adsorbed EVs.

### CLSM analysis of EV-SLB interactions

To gain further insight into EV adsorption, interaction, and organization on the surface of the charged SLBs, we performed CLSM experiments on the same SLB-EV systems analyzed by QCM-D. The experiments focused on three different DOPC/DOTAP SLB configurations as substrates. More precisely, we studied (i) pure DOPC as a negative control for EV adsorption (ii) 40% DOTAP, and (iii) 50% DOTAP, where the latter two represent the SLB compositions for which the QCM-D experiments recorded the most dramatic change in EV adsorption. Fig. S4 displays representative images of the obtained SLBs and confirms the formation of homogeneous, fluorescent lipid bilayers covering the entire surface for all three lipid mixtures.

After a 30-minute equilibration time, 100 μl of EV solution at the same concentration as used in the QCM-D experiments was added to the sample wells and images were collected at specific time intervals. Fig. 3 reports representative top-view 2D images of the SLB-EV systems, collected after 1 hour of incubation after EV injection. The green channel accounts for the fluorescence signal from the SLBs (left column), while the fluorescence from the EVs is displayed in red (centre column). As a result, the superposition of the two channels (right column) in areas where the green and red fluorescence intensities are comparable appears yellow. The results demonstrate that, after one hour of incubation, no EVs had visibly adsorbed on the pure DOPC SLB (top row). Moreover, Movie S7 shows that even those EVs that approach the DOPC SLB surface did not adsorb but were free to move away from the SLB, confirming the negligible adsorption recorded in the respective QCM-D experiment. EVs start to adsorb on the 40% DOTAP SLB, forming very few localized fluorescent aggregates (Fig. S5). On the contrary, EV adsorption on the 50% DOTAP SLB was much higher; after one hour, as shown in Fig. 3 (centre column, bottom row) the 50% DOTAP SLB is characterized by an intense red fluorescence signal across the whole surface, indicating strong adsorption of EVs compared to the limited adsorption on the 40% DOTAP SLB. To ensure that the fluorescence signal is solely due to adsorbed EVs rather than to free dye adsorption, we performed control experiments on the same SLB systems without EVs but with the red dye Cy5 present. As shown in Fig. S6, the red fluorescence signal effectively vanishes in the absence of EVs for all three SLBs, showing that the signal is indeed due to EV adsorption.

**Figure 3:**
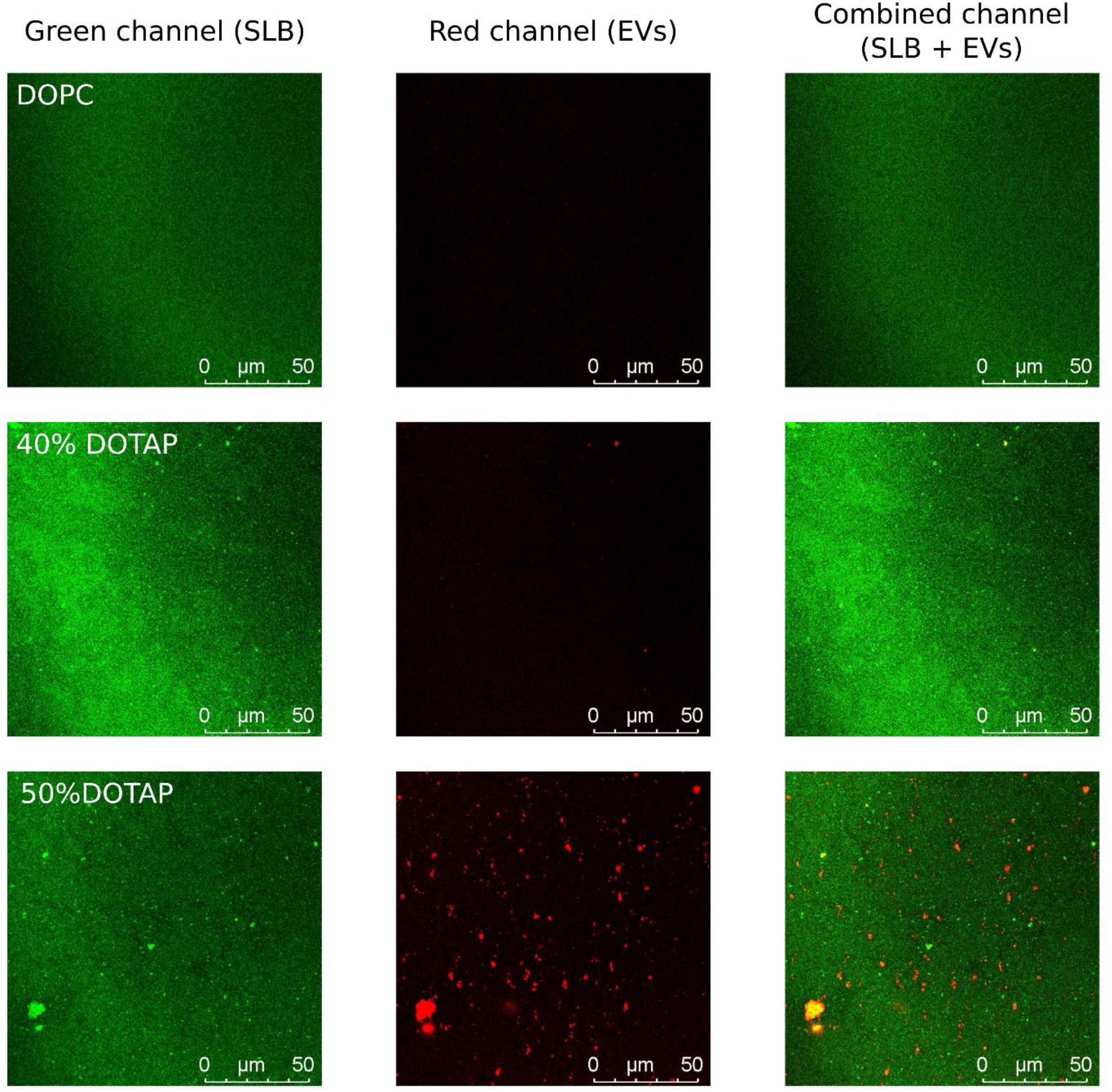
Representative top-view 2D CLSM images of the probed SLB-EV systems: DOPC (top row), 40% DOTAP (middle row) and 50% DOTAP (bottom row), collected 1 hour after EV injection. The green and red channels respectively report the fluorescent intensities of SLBs (left column) and EVs (centre column). The superposition of the two channels is shown in the right column.

The CLSM results thus highlight the strong attractive interaction between EVs and the more highly charged 50% DOTAP SLB and support our interpretation of the QCM-D results. Notably, the combined fluorescence signal from the 50% DOTAP SLB + EVs system (Fig. 3 right column, bottom row) is still mostly saturated by the green fluorescence from the underlying SLB, meaning that the adsorbed EVs did not rupture and form a tightly packed lipid bilayer possessing a comparable intensity. Fig. S5 furthermore shows 3D CLSM images of the adsorbed EVs on the 40% and 50% DOTAP SLBs; in the latter case, the image confirms that EVs adsorb on the entire SLB, even forming large aggregates in specific regions, but without rupturing and merging into a flat lipid bilayer. To address the mechanistic details of the EV-SLB interaction and obtain a more thorough understanding of the colloidal forces driving each single vesicle to adsorption, we will in the next section instead formulate a model for the interaction energy between the SLB and the EVs.

### Theoretical modeling of EV-SLB interaction

To verify that the electrostatic interaction between EVs and oppositely charged SLBs is indeed the dominant driving force for EV adsorption observed in microscopy and QCM-D, we built a semiquantitative description based on the Poisson-Boltzmann (PB) equation describing the electrostatic interactions^39^. Unlike in standard DLVO theory, we consider a full nonlinear PB description of the interaction between *unequally* charged particles, which furthermore makes it necessary to solve the PB equation without any symmetry about the midplane. In addition, we will include the effect of short-ranged, repulsive “steric-hydration” forces that are known to generically act between hydrated lipid bilayers.

Within the PB framework, two oppositely charged macroions or surfaces immersed in a salt solution will attract at long range due to the favorable free energy associated with releasing the counterions into the bulk solution as the two surfaces approach each other^29,39^. If the two surface charge densities *σ*_1_ and *σ*_2_ are equal and opposite (i.e., *σ*_1_ = −*σ*_2_), this attraction will persist monotonically down to close contact, where the two surfaces will neutralize each other completely, releasing all counterions into the bulk. In the typical case where *σ*_1_ ≠ −*σ*_2_, the attraction will instead change into a repulsion at the separation where all “excess” counterions have been released into the bulk; the layer of remaining counterions that neutralize the total charge *σ*_1_ + *σ*_2_ of the two surfaces is then compressed as the surfaces approach further, leading to a repulsive entropic force^29^. Thus, within the PB framework, we expect an electrostatic interaction free energy with a well-defined minimum at intermediate separations, whose depth and location depend on the respective surface charge densities and the salt concentration.

We model a single EV as a negatively charged sphere with constant and homogeneous surface charge density *σ*_EV_, while the SLB is modelled as a rigid flat plane with corresponding charge density *σ*_SLB_. For a monovalent (1:1) salt solution at bulk concentration *c*_0_, the nonlinear PB equation for two flat parallel surfaces reads

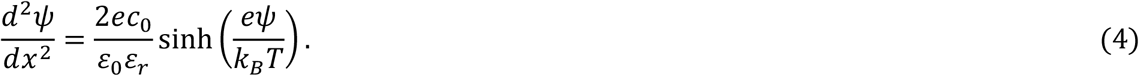

Here, *ψ*(*x*) is the electrostatic potential, *x* ∈ [0, *D*] the position along the surface normal, *ε*_0_ is the vacuum permittivity, *ε*_*r*_ the relative dielectric permittivity of water, and *k*_*B*_*T* the thermal energy. The SLB surface charge density *σ*_SLB_ was calculated using Eq. (3) and listed in Table 1. The EV surface charge density *σ*_EV_ is more difficult to estimate *a priori*; we therefore manually adjusted this value to the one that gave the best agreement with the experimental estimate of the adsorbed amount of EVs for the 50% DOTAP system, as discussed below and shown in Fig. 5, yielding a value of *σ*_EV_ = −0.0094 Cm^-2^. By numerically integrating Eq. (4) as described in Materials and Methods, we obtained the electrostatic free energy *A*_EI_(*D*) between the spherical EV and the flat SLB, where *D* is the minimum separation between the two surfaces. The electrostatic interaction curves are plotted in Fig. 4a, showing that these alone indeed lead to a non-monotonic behavior, with a free energy minimum whose depth is roughly independent of the SLB charge (controlled by the DOTAP fraction) but whose position gradually moves towards larger separations as the DOTAP content is increased.

**Figure 4:**
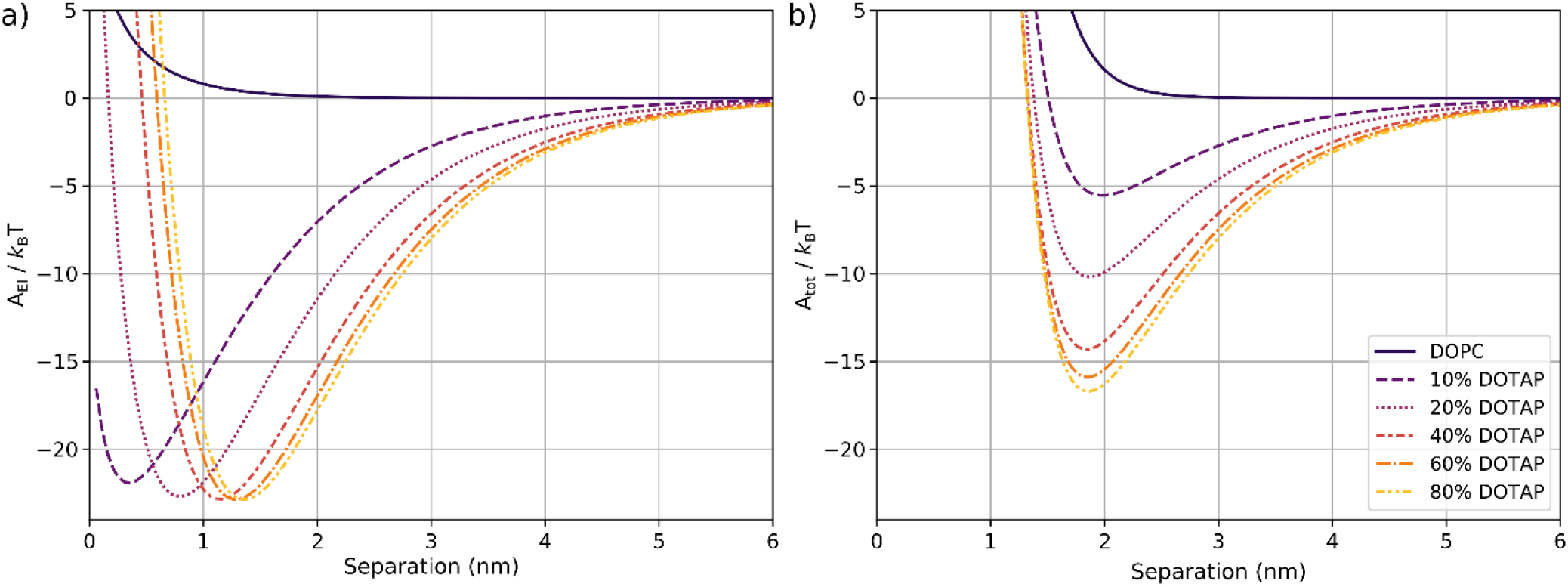
Interaction free energy for different SLB compositions, including the effect of (a) electrostatic forces alone and (b) electrostatic forces and short-ranged steric-hydration forces as given by Eq. (5) with *A*_0 =_ 4500 *k*_*B*_*T* and *D*_0 =_ 0. 25 nm.

**Figure 5:**
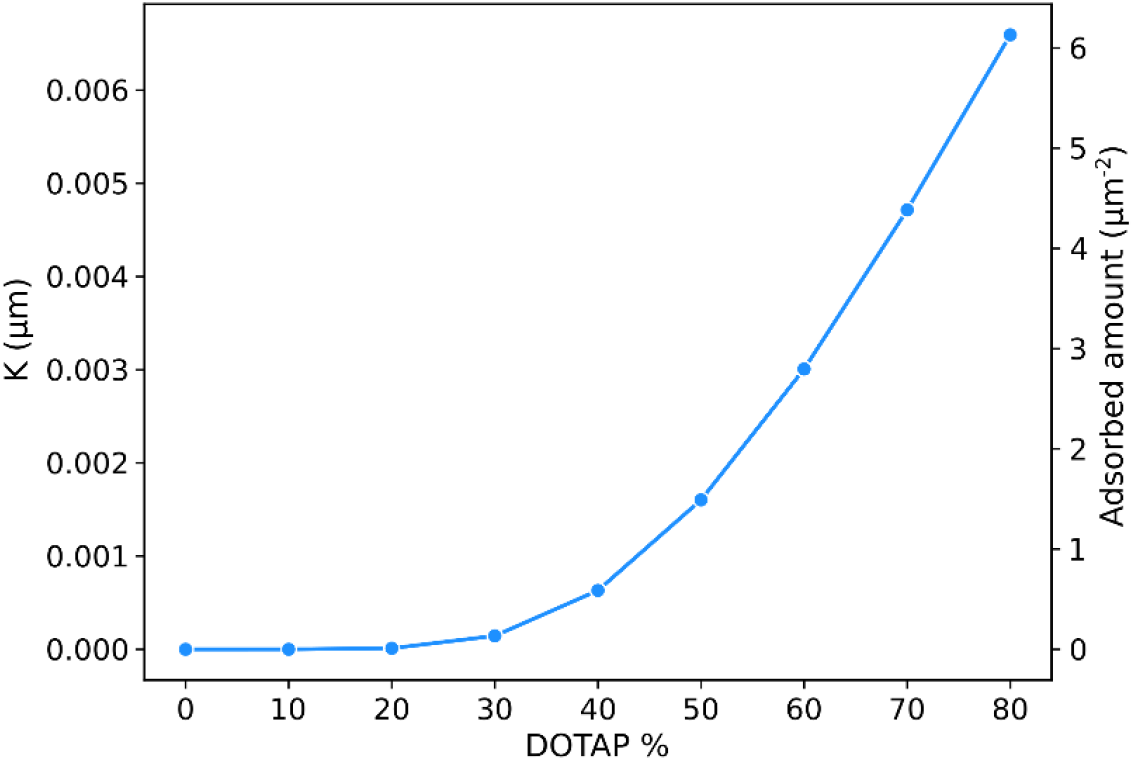
Calculated values of the equilibrium constant *K* for EV adsorption (left ordinate axis) and the adsorbed amount *Γ* (right axis) as a function of the SLB composition, as calculated from Eq. (6) and using the measured bulk concentration *c*_EV =_ 9. 3 × 10^8^ ml^-1^.

For lipid bilayers in aqueous solution, it is well-established that a short-ranged, repulsive force acts to prevent adsorption and adhesion due to attractive van der Waals forces, even in the absence of electrostatic stabilization mechanisms^13^. While the mechanistic origin of this “steric-hydration” force is still debated, it is empirically well characterized as an exponentially decaying interbilayer pressure, *P*_SH_ = *P*_0_exp (−*D*/*D*_0_), with a decay length *D*_0_ on the order of 1 nm. Since we cannot easily measure the force curve between our mixed DOPC/DOTAP bilayers and the complex EV surface, we instead used the parameters *D*_0_ = 0.25 nm and *P*_0_ = 5 × 10^8^ N/m^2^ previously measured for the interaction between uncharged lecithin bilayers^36^. Importantly, this empirically measured short-range force represents the *total* force acting between two SLBs, and thus implicitly also includes the short-range part of van der Waals attractions and any other non-electrostatic interactions acting between the SLBs. Applying the Derjaguin approximation and integrating the pressure twice, we obtain the repulsive interaction between a sphere and a plane:

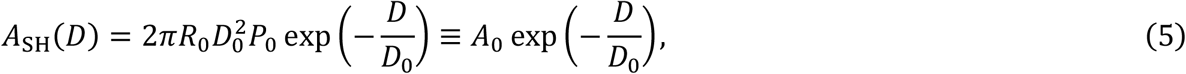

where *R*_0_ is the EV radius and our parameter values yield *A*_0_ = 4500 *k*_*B*_*T*. Due to this large prefactor, the steric-hydration force will be dominant up until a separation of *D* ∼ 3 nm, beyond which an attractive van der Waals force sets in. The long-range part of the van der Waals force is however expected to be small, and we thus neglected it in our description. In Figure 4b, we plotted the total interaction energy *A*_tot_(*D*) = *A*_EI_ + *A*_SH_ for SLBs with DOTAP fractions from 0 to 80%. As can be seen, the combination of the two forces makes the attractive minimum change its depth significantly with SLB charge density, while remaining at a roughly constant separation of *D* ≈ 2 nm. This indicates that we should expect a monotonically increasing EV adsorption with increasing DOTAP fraction.

To further quantify this observation, we note that the equilibrium constant *K* for adsorption of EVs on the SLB surface can be related to the interaction free energy *via*^39^

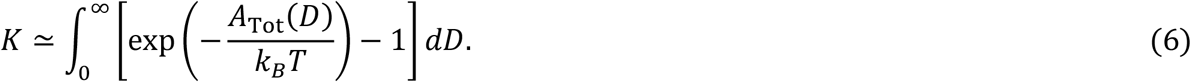

In Fig. 5 (left ordinate axis), we report the calculated values of *K* from Eq. (6) as a function of the SLB composition. The reported values clearly show that our model indeed predicts an electrostatically induced adsorption starting from DOTAP fractions above 30%, in excellent agreement with our experimental results.

The modelling also shows that the adsorbed amount is expected to increase continuously with the DOTAP fraction, in accordance with the observed frequency shifts in QCM-D. This fact is not obvious *a priori*, as a simple picture based on the electrostatic interaction alone would lead to a sharp initial increase in adsorbed amount already for small DOTAP fractions, followed by an essentially constant plateau. This can be seen by the fact that the depth and width of the free energy minimum in the purely electrostatic picture (Fig. 4a) are essentially constant beyond 10% DOTAP, a picture that changes qualitatively when the effect of short-ranged repulsive forces is added (Fig. 4b).

The calculated equilibrium constant *K* can furthermore be related to the number of adsorbed EVs per unit area, Γ, *via* its definition Γ = *Kc*_EV_, where *c*_EV_ ≈ 9.3 × 10^8^ ml^−1^ is the measured bulk EV concentration after adjusting for the different dilutions. We experimentally estimated Γ from the CLSM image at 50% DOTAP (Fig. 3, bottom row, center panel) by measuring the total area of the bright spots in the image, using the software *Gwyddion*^40^. The number of adsorbed EVs per unit area was obtained by dividing the total area by the projected area 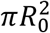 of a single EV and by the total image area, yielding Γ ≈ 1.49 μm^−2^. Obviously, this estimate is highly approximate and potentially also affected by the fact that the bare size of individual EVs is below the CLSM resolution limit (∼250 nm); nevertheless, we believe that it is accurate enough to perform a semiquantitative comparison between the PB modelling and experimental results. The experimentally estimated value of Γ was then used to adjust the unknown EV surface charge density *σ*_EV_ = −0.0094 Cm^-2^. This value corresponds to an EV surface potential of *ψ*_0_ ≈ *σ*_EV_/(*ε*_0_*ε*_*r*_κ) = −10.6 mV, with κ being the inverse Debye screening length, and where the linear relation between surface potential and surface charge is accurate for low potentials such as those measured for EVs. Just as expected, the estimated value of the surface potential has the same sign somewhat larger magnitude than the measured EV *ζ* potential of −77 mV (see Table 1), since the *ζ* potential is measured in the slip plane of the double layer where part of the surface charge has been screened.

The fact that we can obtain an excellent agreement between theoretical modelling and experimental results using reasonable parameter values indicates that EV adsorption is indeed controlled by an intricate interplay between the long-ranged attractive and short-ranged repulsive forces acting in the system. The theoretical results furthermore explain the experimental observation that EVs will start to adsorb onto the SLB above a composition of approximately 40% DOTAP and that they do not reach adhesive contact with the SLB, but rather remain adsorbed at some finite distance outside the membrane. This is in accordance with previous experimental results^36^ indicating that electrostatic interactions are typically not sufficient to induce fusion, likely due to the repulsive barrier induced by compression of the remaining counterions. In this picture, fusion will only occur when deformations within the EV membrane and SLB expose the hydrocarbon groups and hence give rise to attractive hydrophobic interactions.

## Conclusion

The results presented here demonstrate that electrostatic interactions can be used to control the adsorption of EVs onto charged lipid membranes. More precisely, we have shown that EVs readily adsorb, without merging with the SLB, onto membranes possessing opposite surface charge densities *σ* higher than a specific threshold value. When adsorption occurs, its extent is found to be roughly proportional to *σ*. Modelling the system within the framework of nonlinear Poisson-Boltzmann theory confirmed the experimental results and showed that electrostatic interactions are indeed sufficient to cause the irreversible binding of EVs to charged SLBs, but prevent attraction down to molecular contact due to the remaining counterions. This suggests that electrostatic interactions may play a role in vesicle-mediated intercellular communication, since they are capable of causing irreversible binding of EVs with the charged sites of the cell membrane. However, these forces alone are likely not sufficient to account for EV-fusion or uptake, phenomena where other types of interactions come into play. This is in agreement with previous works that documented EV fusion due to either the energetically favorable conditions encountered along the host membrane phase boundaries^26^ or antibody-mediated attractive interactions^27^. Taken together, our findings highlight the fundamental role played by surface charge in the interaction of EVs at the nanoscale, adding to the knowledge about their stability and binding propensity to lipid interfaces. Given the growing interest in employing EVs for numerous biomedical applications, our results are of potential interest for developing efficient EV immobilization strategies necessary for controlling and manipulating systems based on these natural lipid-based nanocarriers.

## Acknowledgements

JS acknowledges enlightening discussions with Håkan Wennerström. We thank Marije Kleinjan (Utrecht University, Department of Biomolecular Health Sciences, The Netherlands) for technical support in preparing milk EV samples and Valentina Moccia (Department of Comparative Biomedicine and Food Science, University of Padua, Italy) for technical support in the NTA characterization. This work has been supported by the European Community through the evFOUNDRY project (H2020-FETopen, ID: 801367).

## Notes

### Competing Interest Statement

The authors have declared no competing interest.

