## Supplementary Information for "Electrostatic interactions control the adsorption of extracellular vesicles onto supported lipid bilayers"

**Table S1: Weight percentages of DOTAP and DOPC used for liposomes with different surface charge. The "Sample name" column indicates the nomenclature used for referring to particular liposome/SLB compositions throughout the manuscript.**

| Sample name | DOTAP % (w/w) | DOPC % (w/w) |
|---|---|---|
| DOPC | 0 | 100 |
| 20% DOTAP | 20 | 80 |
| 30% DOTAP | 30 | 70 |
| 40% DOTAP | 40 | 60 |
| 50% DOTAP | 50 | 50 |
| 75% DOTAP | 75 | 25 |

**Table S2: Values of hydrodynamic radius and respective polydispersity index (PDI) for all the probed liposome dispersions, measured by Dynamic Light Scattering (DLS).**

| Sample name | Hydrodynamic Radius (nm) | PDI |
|---|---|---|
| DOPC | 95.8 ± 0.3 | 0.164 ± 0.004 |
| DOTAP 20% | 77.3 ± 0.7 | 0.186 ± 0.015 |
| DOTAP 30% | 75.8 ± 0.2 | 0.208 ± 0.008 |
| DOTAP 40% | 89.6 ± 0.6 | 0.227 ± 0.007 |
| DOTAP 50% | 76.1 ± 0.2 | 0.125 ± 0.002 |
| DOTAP 75% | 74.9 ± 0.4 | 0.166 ± 0.212 |

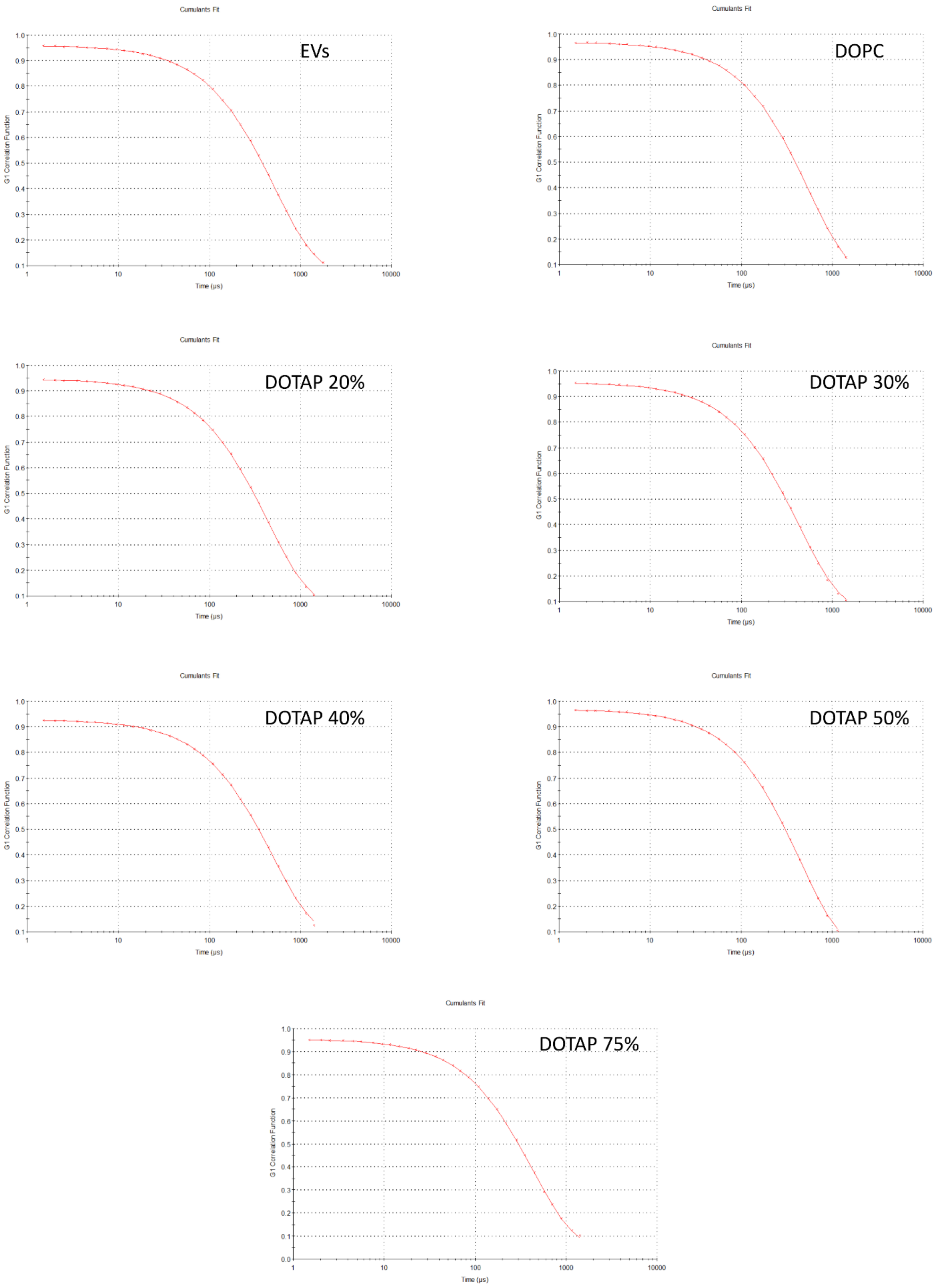


*Figure S1:* **Autocorrelation functions as a function of decay time and the corresponding best fits according to the cumulant algorithm, obtained for the EV samples and for all the probed liposome preparations.**

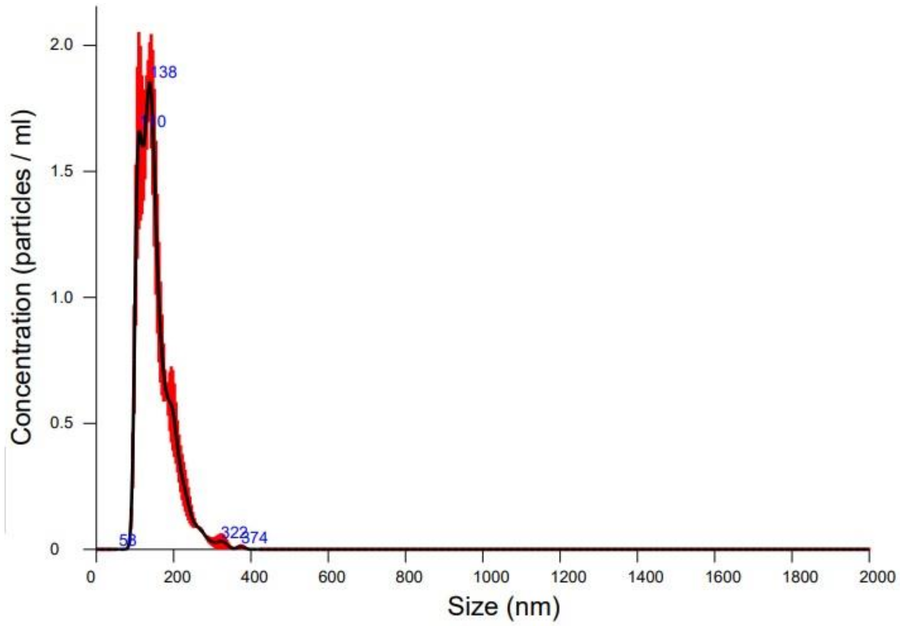


***Figure S2:*** **Nanoparticle Tracking Analysis (NTA) of the EV sample at a 500x dilution in PBS. The red shading shows the standard deivation calculated from the triplicate 60-sec captures. The measured average vesicle diameter was $\mathbf{151.5 \pm 5.5}$ nm.**

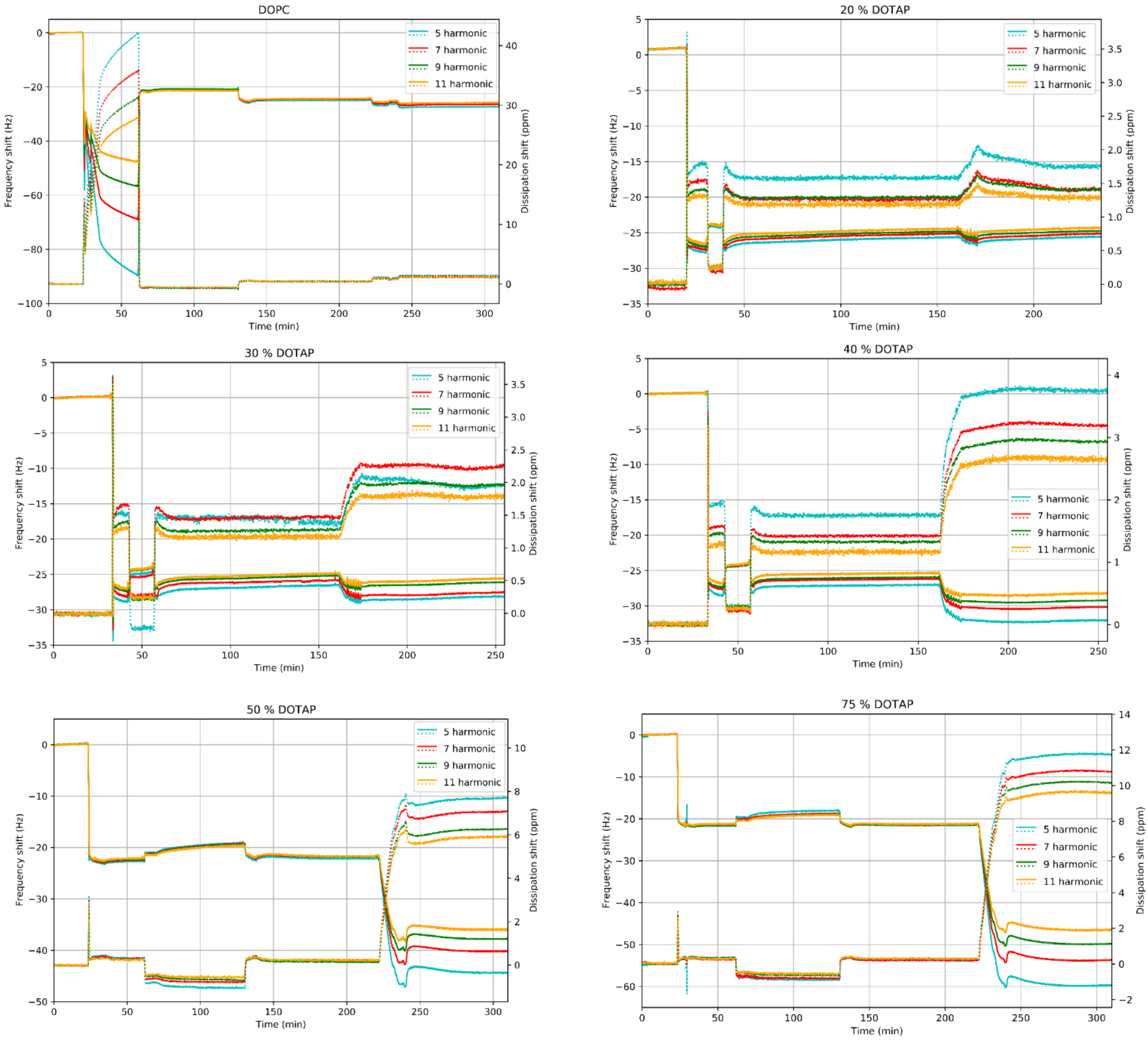


***Figure S3:*** **QCM-D data for all studied SLB compositions. Solid lines indicate the frequency shift values while the dotted lines report the dissipation shifts for the 5th, 7th, 9th and 11th harmonics (cyan, red, green and orange, respectively).**

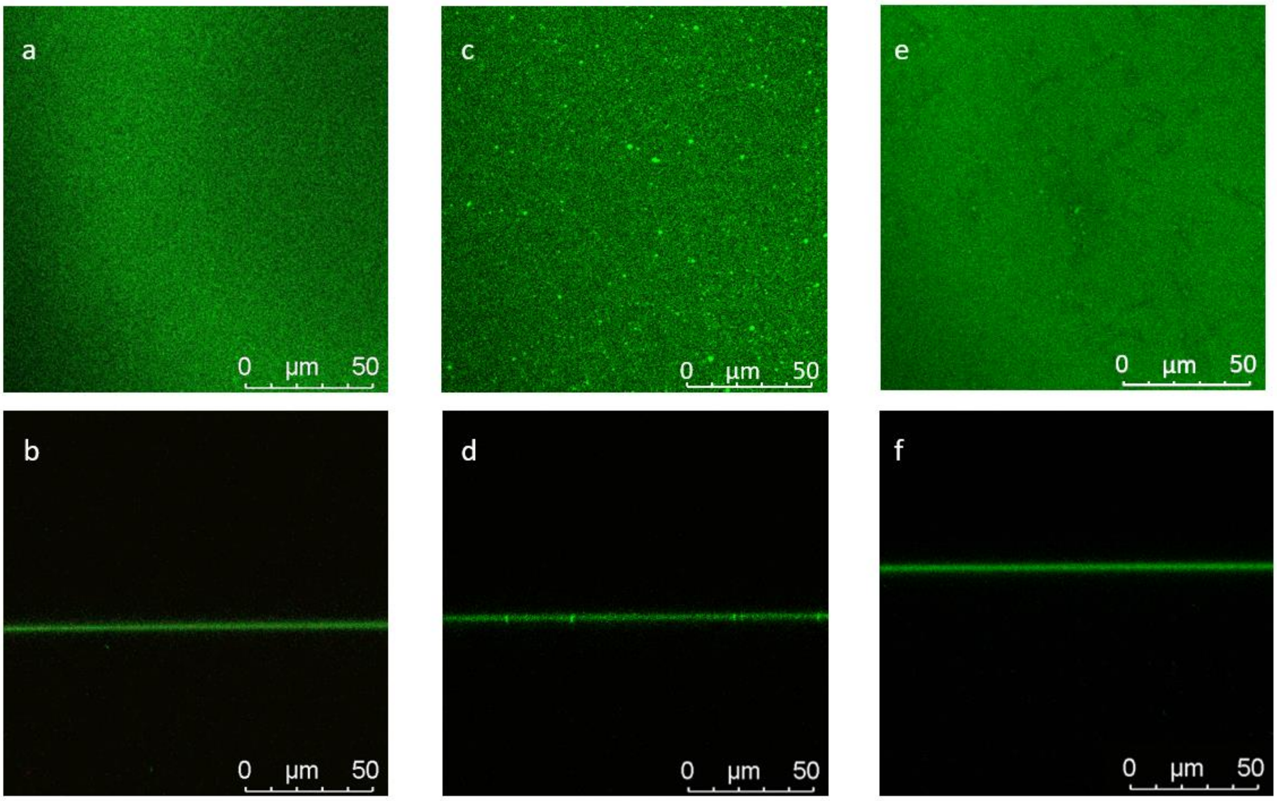


***Figure S4***: **Representative CLSM top- and side-view images of SLBs made from pure DOPC (a,b), 40% DOTAP (c,d), and 50% DOTAP (e,f).**

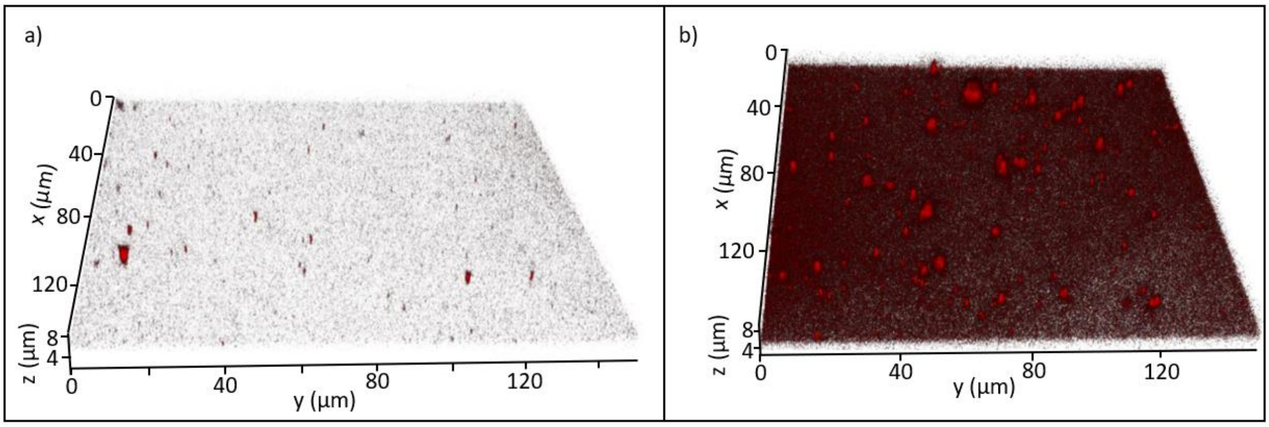


***Figure S5***: **CLSM 3D images showing EVs adsorbed on (a) 40% DOTAP and (b) 50% DOTAP SLBs. The adsorbed EVs cover the entire surface at the higher charge density, including the presence of larger aggregates. In contrast, the adsorption on the 40% DOTAP SLB is dramatically lower and less homogeneous.**

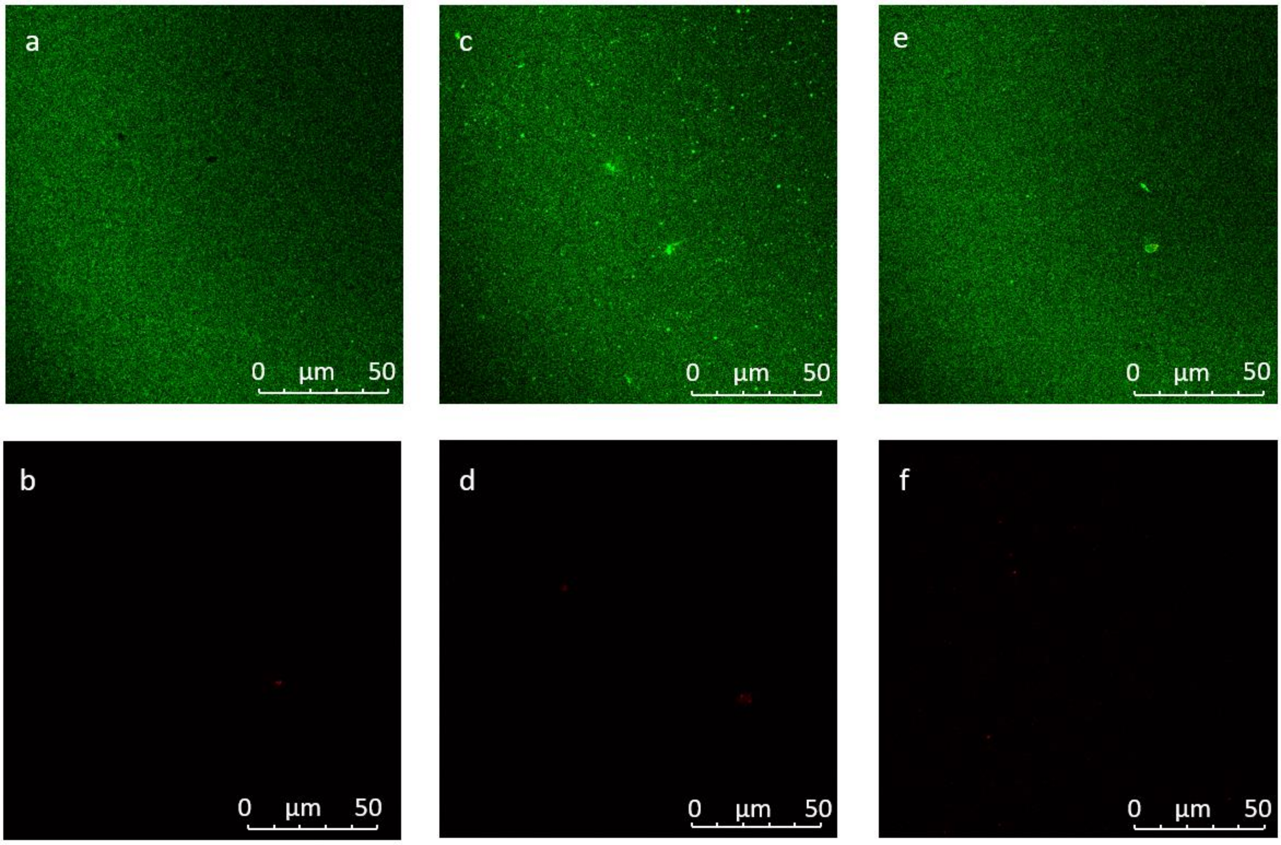


*Figure S6*: **Representative top-view CLSM images of SLBs in PBS collected 2 hours after the injection of 100 µl of Cy5 dye (0.1 mol%). The top and bottom rows respectively show the green and red (Cy5 dye) fluorescent channels for SLBs made from pure DOPC (a,b), 40% DOTAP (c,d), and 50% DOTAP (e,f). The images in the bottom row show that, in the absence of EVs, the fluorescent signal coming from the red dye is negligible for all SLB compositions.**
